# Pkn1 kinase is a critical regulator of cell division in *Chlamydia trachomatis*

**DOI:** 10.64898/2026.09.24.754023

**Authors:** Junghoon Lee, John V. Cox, Scot P. Ouellette

## Abstract

*Chlamydia trachomatis* is an obligate intracellular, developmentally regulated pathogen with a reduced genome as a consequence of adapting to the host cell environment. Despite possessing only eight transcription factors, its developmental cycle is tightly regulated. Hence, we hypothesize that factors like post-transcriptional and post-translational modifications play significant roles in controlling developmental progression. *Chlamydia* encodes serine/threonine kinases, including an ortholog of Pkn1. In this study, we characterized the function of Pkn1 in chlamydial biology and revealed its role as a cell division regulator. We designed an anhydrotetracycline (aTc)-inducible construct encoding Pkn1 tagged with six histidines at the C-terminus (Pkn1_6xH). Wild-type Pkn1 localized primarily at the division septum whereas a kinase-null Pkn1 isoform localized in the cytosol. To further explore the association of Pkn1 with the divisome, we designed a plasmid encoding an aTc-inducible CRISPRi system targeting *pkn1* (*pkn1*KD). Intriguingly, decreased *pkn1* transcript levels resulted in enlarged chlamydial cells with irregular coccoid shapes akin to penicillin-treated cells, indicating a block in cell division when *pkn1* transcripts are reduced. These effects were complemented by expressing an additional wild-type copy, but not a kinase-null copy, of *pkn1* during knockdown. Using a bacterial two hybrid system, we identified an interaction between Pkn1 and MreB, a cytoskeletal protein forming a ring at the septum in *Chlamydia*. We showed the co-localization of these proteins in *Chlamydia* and that inhibition of MreB function disrupts Pkn1 localization. Overall, our data indicate an essential function for Pkn1 activity in regulating cell division in *Chlamydia*.

**Importance:** The Serine/Threonine protein kinase superfamily is present in prokaryotes, and Pkn1 is a member of this superfamily in *Chlamydia trachomatis*. Chlamydial Pkn1 was previously characterized as a S/T protein kinase *in vitro*, but no *in vivo* studies had been performed to characterize its function in *Chlamydia*. In this study, we identified the *in vivo* function of Pkn1 as a regulator of chlamydial cell division. Pkn1 localized to the chlamydial division septum, and disrupting its expression blocked cell division. Our findings are important in understanding the molecular mechanism of chlamydial cell division and contribute to identifying new targeted therapeutic strategies for chlamydial infectious diseases.

## Introduction

*Chlamydia trachomatis* is a developmentally regulated, obligate intracellular bacterium that replicates within its host cell inside of a membrane-bound niche termed an inclusion (1). Its developmental cycle alternates between two morphologically and functionally distinct cell types: the elementary body (EB), which is infectious but non-dividing, and the reticulate body (RB), which is non-infectious but dividing (See ref.(2)). Transition between these two forms is a hallmark of the chlamydial developmental cycle. During the replicative phase, RBs divide by a polarized MreB-dependent cell division mechanism rather than by canonical FtsZ-dependent binary fission(3). This process is coordinated by the DNA translocase FtsK, which recruits divisome components to a distinct site on the inner membrane of the RB (3, 4). During division, the actin-like protein MreB forms a ring structure at the septum by detecting membrane curvature produced by cardiolipin, and MreB function is required for peptidoglycan (PG) ring formation (3, 5–7). Although this unusual division process is unique and interesting, the regulatory mechanisms governing its spatial and temporal control remain poorly defined.

Protein phosphorylation is one of the most widespread and fundamental mechanisms of signal transduction across all domains of life(8). This reversible post-translational modification enables cells to rapidly sense and respond to changes in their environment, metabolic state, and physiological conditions, thereby supporting cellular adaptation, homeostasis, and survival. Phosphorylation can regulate a broad range of cellular processes by altering the enzymatic activity, conformation, stability, subcellular localization, or interaction partners of target proteins(9). It can also influence gene expression by modulating the activity of transcription factors, RNA polymerase-associated proteins, and other components of transcriptional regulatory pathways(10). The phosphorylation state of cellular proteins is dynamically controlled by the opposing activities of protein kinases, which transfer phosphate groups—typically from ATP—to specific amino acid residues, and protein phosphatases, which remove these phosphate groups(9, 11). The coordinated actions of these enzymes allow phosphorylation-dependent signaling pathways to function as precise and reversible molecular switches that integrate cellular signals and produce appropriate physiological responses.

Bacterial phosphorylation systems can modulate gene expression, alter protein conformation, control enzymatic activity, and reorganize cellular architecture, thereby promoting adaptation and survival under changing conditions. Among bacterial protein kinases, the serine/threonine protein kinase (STK) superfamily represents a conserved class of enzymes that catalyze transfer of the γ-phosphate of ATP to a serine or threonine hydroxyl group on a target protein. Because these enzymes share structural and mechanistic similarity with eukaryotic Hanks-type kinases, they were originally described as eukaryotic-like protein kinases. However, the first bacterial S/T protein kinase was identified from *M. xanthus* in 1991 and characterized to function in the development of *M. xanthus*(12). Subsequent genomic and biochemical studies demonstrated that STKs are widely distributed in bacteria and regulate diverse processes, including metabolism, stress responses, virulence, and cell division(13).

*C. trachomatis* encodes two genes, *pkn1* and *pknD*(14), and the proteins encoded by these genes possess the conserved domain of the STK superfamily. A prior study recombinantly expressed and purified Pkn1 and PknD from *E. coli* and confirmed their S/T protein kinase activity by *in vitro* kinase assays(15). In addition to the kinase activity, the K42 residue of Pkn1, which is a predicted ATP-binding residue, was shown to be crucial for Pkn1’s kinase activity(15). However, *in vivo* functions of these proteins in *C. trachomatis* were unknown prior to this study. We hypothesized that post-translational modifications mediated by one or both of these S/T kinases are critical for chlamydial developmental progression. In this study, we characterized Pkn1 using both overexpression and knockdown strategies. Localization to the septum identified Pkn1 as a chlamydial cell division protein. This was supported by the effects of *pkn1* knockdown, which blocked cell division, and interaction of Pkn1 with divisome components. Together, these findings identify Pkn1 as a previously unrecognized regulator of polarized cell division in *C. trachomatis*.

## Results

### Pkn1 localizes at the septum in *C. trachomatis*

Chlamydial Pkn1 contains a conserved STK domain at its N-terminus, including the conserved ATP binding residue K42 as previously reported (Fig. 1A)(15). InterPro analysis predicted that the C-terminal domain is related to sulfatase modifying factor 1(16). Based on this bioinformatics analysis, we generated anhydrotetracycline (aTc) and theophylline-inducible constructs encoding either wild-type Pkn1 or a K42G mutant, in which the conserved ATP-binding lysine was replaced with glycine. Both proteins were tagged with six histidines at the C-terminus (Pkn1_6xH and Pkn1(K42G)_6xH, respectively). Following transformation of these plasmids into plasmid-free chlamydial cells, we infected these transformants into McCoy cell monolayers. The constructs were induced at 4 hours post-infection (hpi) with 5 nM aTc and 0.5 mM theophylline. To examine Pkn1 localization during the first round of division, infected cells were fixed at 10.5hpi and permeabilized prior to antibody labelling. Pkn1_6xH prominently localized to the division septum, where it formed a ring structure resembling that previously reported for chlamydial MreB (Fig. 1B and S1A)(5). We also observed the same localization pattern of endogenous Pkn1, which was detected by Pkn1-specific antibody (Fig. S1B). In contrast, Pkn1(K42G)_6xH rarely localized to the septum and instead exhibited predominantly diffuse cytosolic or punctate localization patterns (Fig. 1B). Quantitative analysis revealed that 71% of cells expressing Pkn1_6xH exhibited septal localization, whereas septal localization was observed in only 6% of cells expressing Pkn1(K42G)_6xH (Fig. 1C). Similar localization patterns for the wild-type and K42G proteins were also observed at 24 hpi (Fig. S1C). Notably, overexpression of either isoform had a limited impact on chlamydial EB production as measured by inclusion forming units (IFUs; Fig. S1D). Together, these findings demonstrate that Pkn1 localizes to the division septum and that its kinase activity is required for proper localization. Collectively, these findings led us to hypothesize that Pkn1 serves an important function in chlamydial cell division.

**Figure 1.**
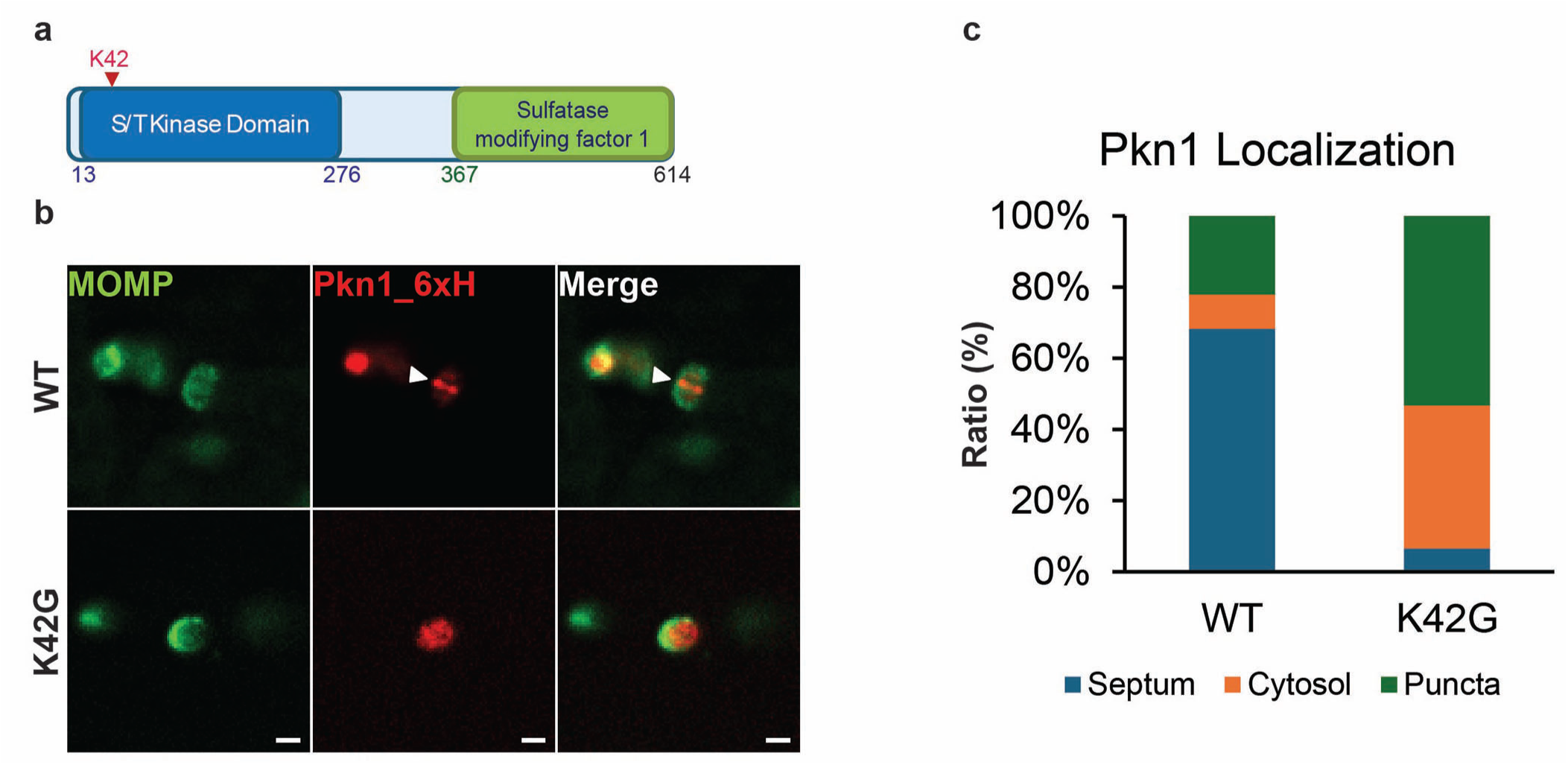
Pkn1 is a S/T protein kinase in *C. trachomatis*. **A.** Scheme of Pkn1’s conserved domains. N-terminal S/T kinase domain was characterized in 2003(15). For C-terminal domain, we predicted the domain with InterPro (https://www.ebi.ac.uk/interpro)(16). **B.** Pkn1 localization at the first dividing cells. Transformant encoding Pkn1_6xH was infected into McCoy cells, and the construct was induced at 4 hpi. The infected monolayer was fixed at 10.5 hpi with fixing solution containing 3.2% formaldenyde and 0.022% glutaraldehyde in 1X PBS for 2 min. And the samples were permeabilized with 90% MeOH for 1 min. MOMP and Pkn1_6xH were stained by indirect immunofluorescence assay (IFA) with Rabbit anti-six histidine and Goat anti-MOMP. **C.** The quantified data of the localization patterns of wildtype and K42G Pkn1 in the first dividing cells. We counted the cells representing septum, cytosol, or puncta localization of wildtype and K42G Pkn1_6xH. 104 cells for wildtype and 92 cells for K42G mutant were counted. IFA images were acquired on a Zeiss AxioImager.Z2 equipped with an Apotome2 using a 100X lens objective. Scale bar: 1 µm.

### *pkn1* knockdown results in abnormal cell morphology

We next made an inducible CRISPR-dCas12 system to generate a conditional knockdown strain targeting the *pkn1* promoter (*pkn1*KD)(17). The *pkn1* gene is the first gene in an apparent operon with *dnlJ*, encoding DNA ligase. The *pkn1*KD strain was compared to a control empty vector (EV) strain encoding only the inducible dCas12 protein. We validated the knockdown system using indirect immunofluorescence assay (IFA) and quantitative PCR (qPCR) (Fig. 2A & D). We confirmed that dCas12 was induced in both strains (Fig. 2A).

**Figure 2.**
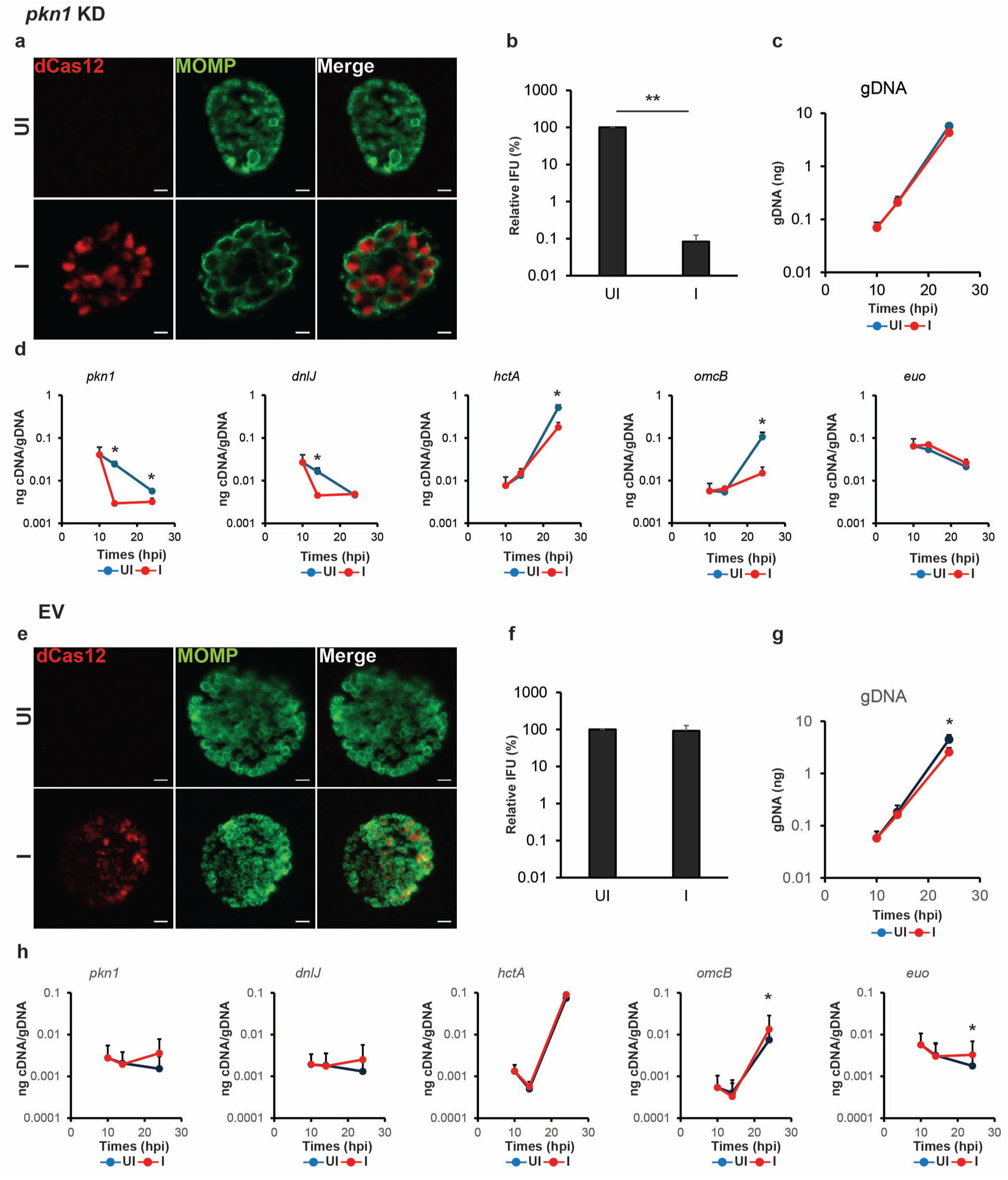
CRISPRi-mediated *pkn1* knockdown shows abnormal cell morphology and lower level of EB progeny production. **(A-D)** McCoy cells were infected with *C. trachomatis* transformed with a construct encoding an anhydrotetracycline (aTc)-inducible CRISPR-dCas12 system targeting the *pkn1* promoter (*pkn1*KD). At 10 hpi, CRISPRi system was induced or not, and DNA and RNA samples were collected at 10, 14, and 24 hpi. Immunofluorescence analysis (IFA) and inclusion forming units (IFU) samples were collected at 24 hpi. **A.** IFA images of the *pkn1*KD strain at 24 hpi. Shown are individual panels for dCas12 and major outer membrane protein (MOMP) labeling as well as the merged image. **B.** Quantification of IFUs from uninduced and induced samples at 24 hpi. **C.** Quantification of genomic DNA by qPCR in uninduced and induced samples. **D.** Quantification of transcripts by RT-qPCR for *pkn1, dnlJ*, *euo*, *hctA*, and *omcB* from uninduced and induced samples. **(E-H)** McCoy cells were infected with *C. trachomatis* transformed with a plasmid encoding an aTc-inducible CRISPR-dCas12 system without target gene (EV). At 10 hpi, knockdown was induced or not, and DNA and RNA samples were collected at 10, 14, and 24 hpi. IFA and IFU samples were collected at 24 hpi. **E.** IFA images of EV strain at 24 hpi. **F.** Quantification of IFU from uninduced and induced samples at 24 hpi. **G.** Quantification of genomic DNA by qPCR in uninduced and induced samples. **H.** Quantification of transcripts by RT-qPCR for *pkn1, dnlJ*, *euo*, *hctA*, and *omcB* from uninduced and induced samples. IFA images were acquired on a Zeiss AxioImager.Z2 equipped with an Apotome2 using a 100X lens objective. Scale bar: 2 µm. UI = uninduced (i.e., -aTc); I = induced (i.e., +aTc) for all sample types. *: p<0.05; **: p<0.001 via two-tailed paired Student’s t-test.

Notably, induction of dCas12 in the *pkn1*KD strain resulted in a pronounced increase in chlamydial cell size compared with the uninduced control (Fig. 2A). Quantitative analysis showed an approximately twofold increase in cell diameter following *pkn1* knockdown (Fig. S2). This enlarged-cell phenotype resembled that observed when chlamydial cell division is arrested by treatment with β-lactam antibiotics (18), suggesting that reduced expression of Pkn1 disrupts normal division (Fig. S3). Reduction of Pkn1 expression caused an approximately three-log decrease in infectious EB progeny production (Fig. 2B), consistent with this. In contrast, chlamydial genomic DNA levels were comparable between the induced and uninduced conditions (Fig. 2C). These results indicate that low levels of Pkn1 strongly impaired cell division.

To confirm knockdown, we measured the transcript levels of *pkn1*, *dnlJ*, encoded 3’ to *pkn1*, *euo* (expressed early in the developmental cycle(19)), *hctA*, and *omcB*. The latter two genes are associated with late stages of the developmental cycle and EB production(19, 20). Induction of dCas12 significantly reduced both *pkn1* and *dnlJ* transcript levels relative to the uninduced condition (Fig. 2D). This decrease was consistent with a polar effect resulting from CRISPR interference targeting the *pkn1* promoter and supports that both genes are encoded in an operon(14). Transcripts for *hctA* and *omcB* were significantly reduced at 24hpi consistent with reduced EB production. No biologically meaningful difference was observed for *euo* transcripts after inducing *pkn1* knockdown.

To determine whether these phenotypes resulted from dCas12 expression alone, we performed the same experiments using an empty vector (EV) CRISPR-dCas12 control construct lacking a targeting crRNA. Induction of dCas12 in this control strain did not produce detectable changes in chlamydial morphology, EB progeny production, genomic DNA levels, or transcript levels (Fig. 2E-H). Thus, the phenotypes observed in the *pkn1*KD strain were dependent on CRISPR-dCas12 system targeting *pkn1* rather than nonspecific effects associated with dCas12 induction.

Collectively, these findings demonstrate that reduced Pkn1 and/or DnlJ is associated with increased chlamydial cell size and a substantial reduction in EB progeny production. Together with the septal localization of Pkn1, these observations support an important role for Pkn1 in the regulation of chlamydial cell division and developmental progression.

### Ectopically expressed Pkn1_6xH complements the *pkn1*KD cell morphology

Because *dnlJ* transcript levels were also decreased by induction of the CRISPR-dCas12 system targeting the *pkn1* promoter region, we sought to determine whether the observed phenotypes resulted from knockdown of *dnlJ* transcription. To address this possibility, we designed a construct to complement *pkn1*KD with expression of DnlJ tagged with six histidines (DnlJ_6xH) and transformed it into chlamydial cells. Although DnlJ_6xH was induced, *pkn1*KD phenotypes were not complemented: cell morphology and EB progeny production were not restored by ectopic expression of DnlJ_6xH (Fig. S4). These findings indicate that reduced DnlJ expression is not sufficient to explain the cell division phenotypes of *pkn1*KD.

We next designed another complementation construct in which Pkn1_6xH was expressed ectopically in the *pkn1*KD background. When Pkn1_6xH was induced, normal chlamydial cell morphology was restored (Fig. 3A), demonstrating that the enlarged cell phenotype resulted from reduced Pkn1 expression. EB progeny production was also increased by approximately two log compared to that of *pkn1*KD. However, EB levels remained approximately one log lower than that of the uninduced control (Fig. 3B). This result suggests that ectopic Pkn1 expression fully rescued the morphological defect but only partially restored EB progeny production of *pkn1*KD. The incomplete recovery of progeny production may reflect the continued reduction of *dnlJ* expression caused by CRISPR interference. This is supported by the fact that complementation of *pkn1* knockdown with both *pkn1* and *dnlJ* completely restored EB production (Fig. S4).

**Figure 3.**
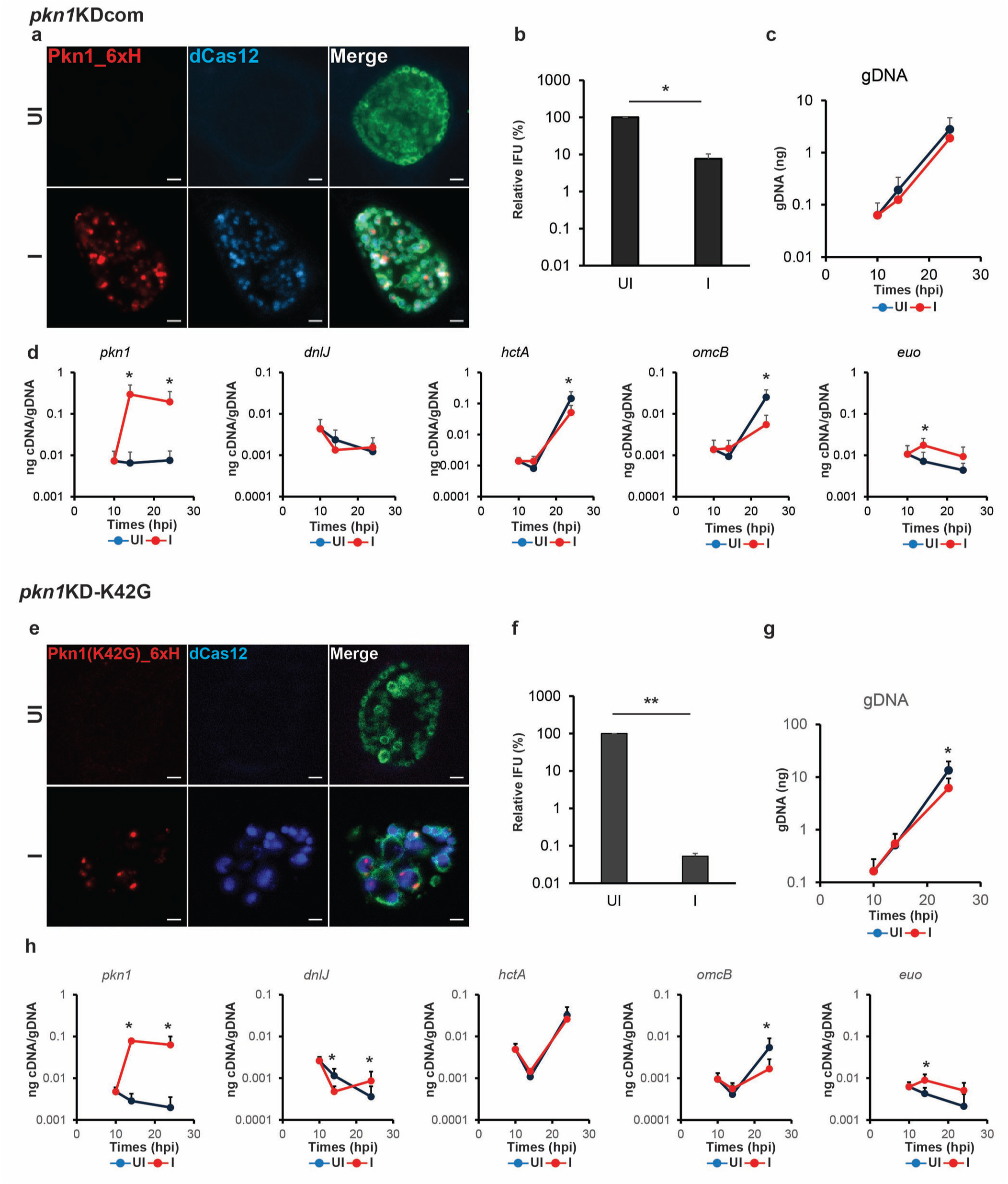
Ectopic expressed Pkn1_6xH complements *pkn1*KD phenotypes. **(A-D)** McCoy cells were infected with *C. trachomatis* transformed with an aTc-inducible plasmid encoding CRISPRi system targeting *pkn1* promoter and Pkn1_6xH (*pkn1*KDcom). At 10 hpi, expression of the construct was induced or not. DNA and RNA samples were collected at 10, 14, and 24 hpi. IFA and IFU samples were prepared at 24 hpi. **A.** IFA images of the *pkn1*KDcom strain at 24 hpi. Shown are individual panels for Pkn1_6xH and dCas12 labeling as well as the merged image with MOMP. **B.** Quantification of IFUs from uninduced and induced samples at 24 hpi. **C.** Quantification of genomic DNA by qPCR in uninduced and induced samples. **D.** Quantification of transcripts by RT-qPCR for *pkn1, dnlJ, euo, hctA, and omcB*. **(E-H)** McCoy cells were infected with *C. trachomatis* transformed with an aTc-inducible plasmid encoding CRISPRi system targeting *pkn1* promoter and Pkn1(K42G)_6xH (*pkn1*KD-K42G). At 10 hpi, expression of the construct was induced or not, and DNA and RNA samples were collected at 10, 14, and 24 hpi. IFA and IFU samples were collected at 24 hpi. **E.** IFA images of *pkn1*KD-K42G strain at 24 hpi. Shown are individual panels for Pkn1(K42G)_6xH, dCas12 as well as the merged image with MOMP. **F.** Quantification of IFUs from uninduced and induced samples at 24 hpi. **G.** Quantification of genomic DNA by qPCR in uninduced and induced samples. **H.** Quantification of transcripts by RT-qPCR for *pkn1, dnlJ, euo, hctA, and omcB* from uninduced and induced samples. IFA images were acquired on a Zeiss AxioImager.Z2 equipped with an Apotome2 using a 100X lens objective. Scale bar: 2 µm. UI = uninduced (i.e., -aTc); I = induced (i.e., +aTc) for all sample types. *: p<0.05 via two-tailed paired Student’s t-test.

To check whether Pkn1’s kinase activity is necessary for complementation of the cell division defect during *pkn1* knockdown, we generated an additional complementation construct expressing Pkn1(K42G)_6xH in the *pkn1*KD background. The K42G substitution disrupts the conserved ATP-binding site and is therefore predicted to abolish or substantially impair Pkn1 kinase activity. Although Pkn1(K42G)_6xH was induced, it failed to complement the *pkn1*KD phenotypes. Chlamydial cells retained the abnormal enlarged morphology, and infectious progeny production remained approximately threefold lower than that of the uninduced control (Fig. 3E-H). These findings demonstrate that the conserved ATP-binding residue K42 is required for Pkn1 function and support the conclusion that Pkn1 kinase activity is critical for normal chlamydial cell division and efficient production of EB progeny.

### Peptidoglycan levels are reduced during *pkn1* knockdown

As described above, *pkn1*KD organisms exhibited an enlarged and irregular morphology resembling that observed following treatment with β-lactam antibiotics, which inhibit PG synthesis (Fig. S3). This phenotypic similarity led us to hypothesize that Pkn1 may contribute to the regulation of chlamydial PG synthesis. In *C. trachomatis*, PG forms a ring-like structure at the division septum(21). To examine PG synthesis and localization, we metabolically labeled chlamydial PG with EDA-DA, followed by detection using Click chemistry(21). We then compared PG labeling in control, *pkn1*KD, and *pkn1*KD complemented organisms (Fig. 4). Under uninduced control conditions, EDA-DA labeling revealed the expected septal ring. In contrast, *pkn1*KD organisms displayed substantially reduced PG-associated fluorescence after knocking down *pkn1* transcripts, and the remaining signal appeared punctate or diffusely distributed along the cell envelope rather than forming a distinct ring. Ectopic expression of Pkn1_6xH in the *pkn1*KD complementation background restored both the intensity of EDA-DA labeling and the characteristic septal PG ring. These findings indicate that Pkn1 is required for proper PG synthesis, incorporation, or organization at the division septum. Together with the enlarged-cell phenotype caused by *pkn1* knockdown, these results support a role for Pkn1 in coordinating septal PG assembly during chlamydial cell division.

**Figure 4.**
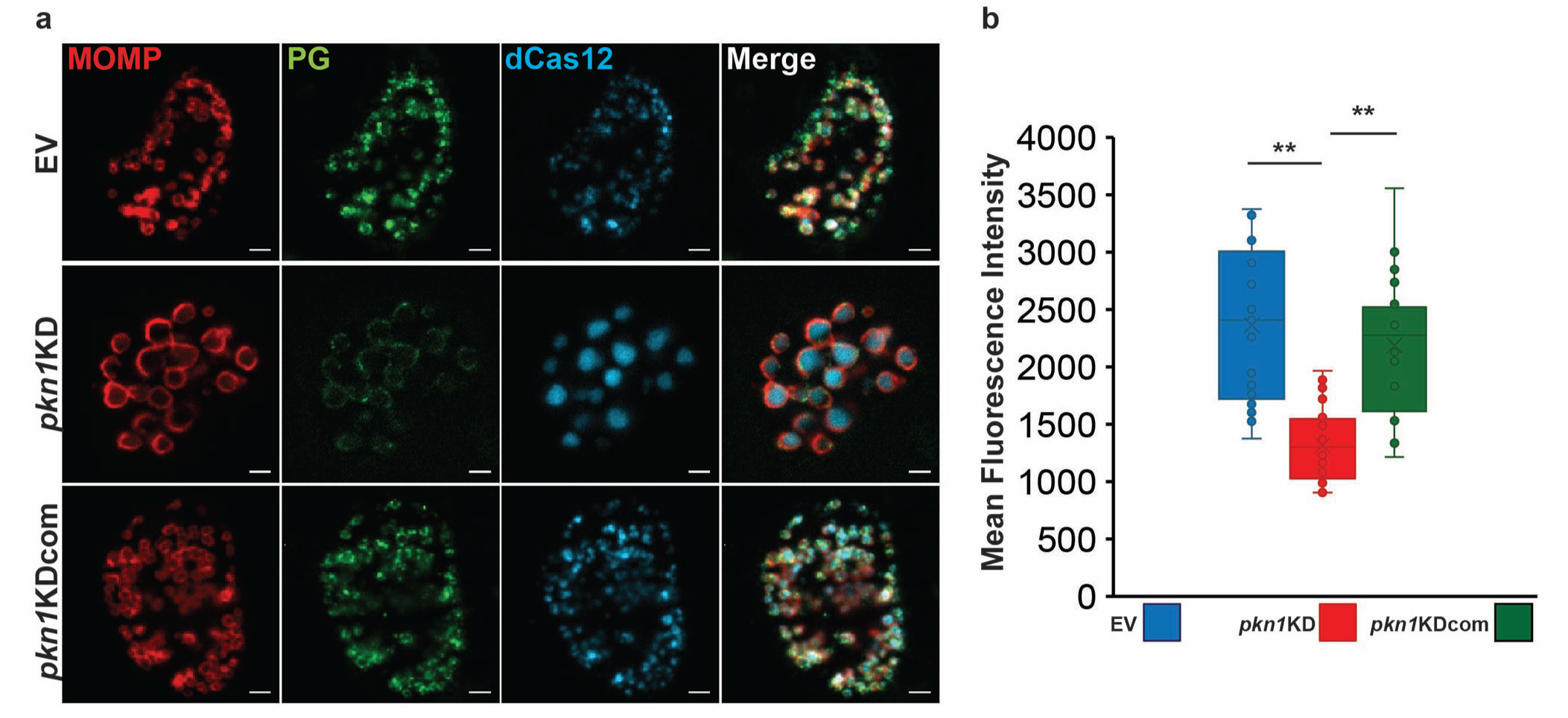
Pkn1 affects the PG synthesis in *C. trachomatis*. **A.** McCoy cells were infected with control strain, *pkn1*KD, or *pkn1*KDcom. The constructs were induced at 10 hpi, and the infected monolayer was fixed with 100% MeOH for 5 min and permeabilized with 0.5% Triton X-100 for 5 min. PG was stained through click chemistry(21). **B.** The quantification of PG levels. The fluorescence intensity of PG was normalized with the area of inclusion by using Fiji software. We measured 21, 26, and 24 inclusions of control, *pkn1*KD, and *pkn1*KDcom, respectively. **: p<0.001 via two-tailed paired Student’s t-test.

### MreB is crucial for optimal localization of Pkn1 at the septum

Because Pkn1 influences septal PG synthesis and organization, we hypothesized that it may interact with proteins of the chlamydial cell division machinery. To identify potential interaction partners, we performed a bacterial adenylate cyclase two-hybrid (BACTH) assay against a panel of chlamydial cell division proteins(22). This analysis revealed an interaction between Pkn1 and MreB (Fig. 5A), a key component of the chlamydial division machinery that coordinates septal PG synthesis. Supporting the BACTH results, we observed the co-localization of Pkn1 and MreB (Fig. 5B). When we quantified the co-localization by using Pearson’s coefficient(23), the value measured was 0.918 ± 0.029, indicating near perfect co-localization. Because MreB forms a ring-like structure at the division septum, we next investigated whether septal localization of Pkn1 depends on MreB organization. Infected cells expressing Pkn1 were treated with A22, a small-molecule inhibitor that disrupts MreB polymerization(6, 24), and Pkn1 localization was subsequently examined. Whereas Pkn1 exhibited prominent septal localization under untreated conditions, A22 treatment caused Pkn1 to become predominantly diffuse within the cytoplasm (Fig. 5C and D). These findings indicate that proper septal localization of Pkn1 depends on polymerized MreB.

**Figure 5.**
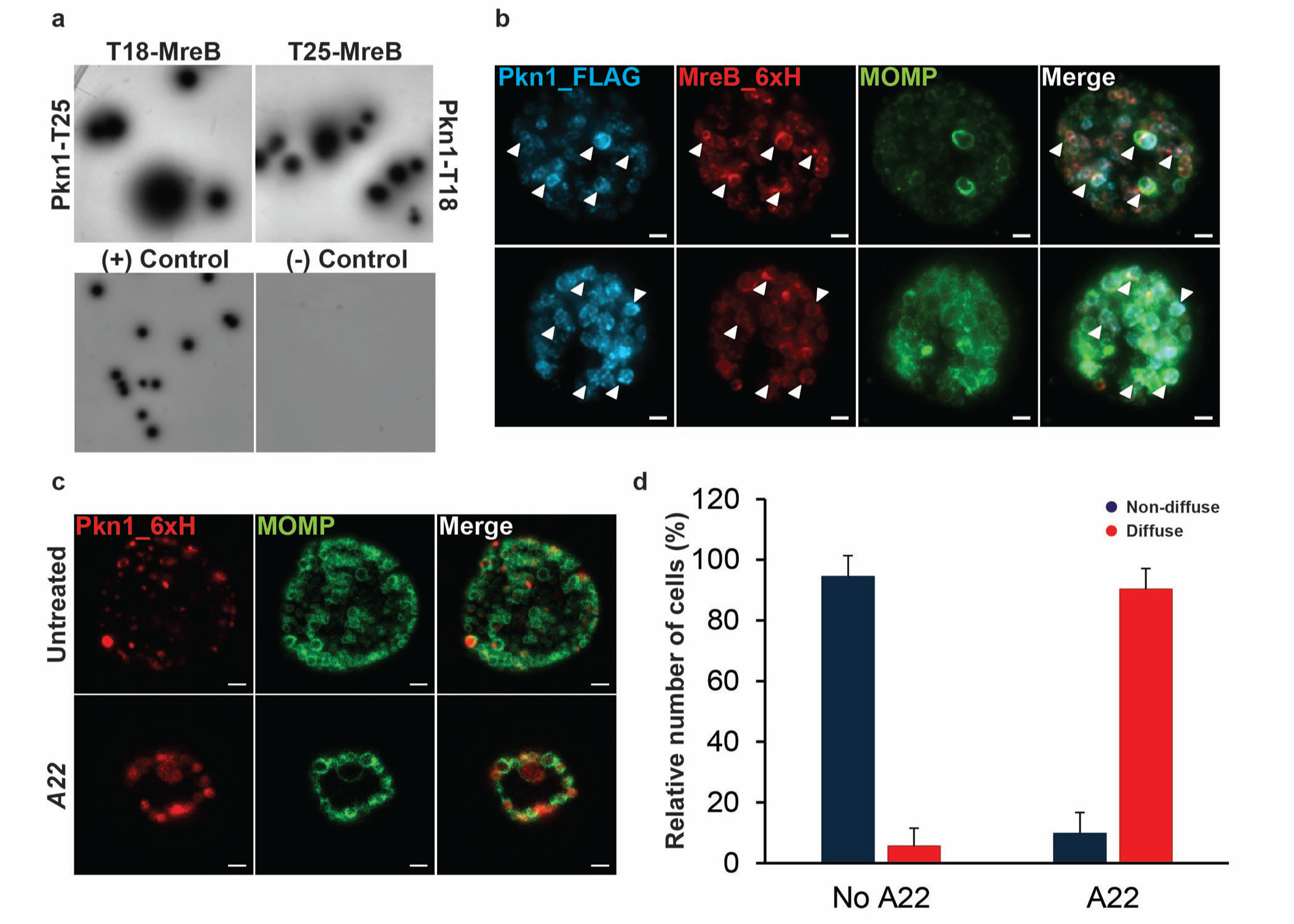
Pkn1 interacts with MreB, and MreB is crucial for Pkn1’s septum localization. **A.** Bacterial adenylate cyclase two hybrid (BACTH) assay. The procedure is described in Materials and Methods. **B.** Co-localization of MreB and Pkn1. The transformant expressing MreB tagged with six histidine and Pkn1 tagged with FLAG was infected into McCoy. At 20 hpi, the construct was induced, and the infected cells were fixed with 100% MeOH for 10 min. The Pearson coefficients were calculated by JACoP in Fiji software, and the value is 0.918 ± 0.029. We measured 66 inclusions’ values. **C.** Pkn1 localization in A22 treated cells. McCoy cells were infected with chlamydial transformant carrying an aTc-inducible construct encoding Pkn1_6xH. At 20 hpi, the construct was induced, and then 75 µM A22 was treated at 22 hpi. The samples were fixed at 24 hpi with 100% MeOH for 10 min. **D.** The number of cells representing diffused Pkn1 in A22. McCoy cells were infected with *C. trachomatis* transformed with an aTc-inducible plasmid encoding Pkn1_6xH. At 10 hpi, expression of the construct was induced, and the 75 μM A22 was treated or not at 22 hpi. The infected monolayer was fixed at 25 hpi with 100% MeOH for 10 minutes, and Pkn1_6xH and MOMP were stained. We counted 269 cells and 332 cells of non-treated and A22-treated samples, respectively. IFA images were acquired on a Zeiss AxioImager.Z2 equipped with an Apotome2 using a 100X lens objective. Scale bar: 2 µm.

## Discussion

Hanks-type STKs were initially associated with eukaryotic signal transduction; however, their presence in bacteria was unambiguously demonstrated in *Myxococcus xanthus* in 1991(12, 25). Subsequent studies have shown that Hanks-type STKs are widely distributed among diverse bacterial species and regulate numerous physiological processes, including cell division, PG homeostasis, metabolism, developmental transitions, and responses to environmental stress(13). In the present study, we characterized the physiological function of Pkn1, one of the two predicted Hanks-type STKs encoded by *C. trachomatis*. Chlamydial Pkn1 was initially characterized by Verma and Maurelli in 2003, who demonstrated that recombinant GST-tagged Pkn1 possessed autophosphorylation activity *in vitro* and could phosphorylate the chlamydial inclusion membrane protein IncG and the other STK protein PknD(15). However, because that study was conducted primarily using recombinant proteins and *in vitro* assays, the physiological function of Pkn1 during the chlamydial developmental cycle remained unknown. This is the first report characterizing the *in vivo* function of Pkn1 in *C. trachomatis*.

Based on the previously reported association between Pkn1 and IncG, we expected that chlamydial Pkn1 might regulate inclusion stability or signaling between *C. trachomatis* and its host cell – including the possibility that Pkn1 may be secreted into the host cell. Unexpectedly, IFA revealed that Pkn1 was localized at the division septum and formed a ring structure during cell division (Fig. 1B and S1A). This localization pattern suggested that Pkn1 functions as a component or regulator of the cell division machinery. Septal localization has also been reported for StkP, the Hanks-type STK of *Streptococcus pneumoniae* (See ref.(26)). StkP is a transmembrane kinase containing an N-terminal cytoplasmic kinase domain and multiple extracellular C-terminal penicillin-binding protein and S/T kinase-associated (PASTA) domains. It is located at the septum during cell division and regulates pneumococcal morphogenesis and cell-wall synthesis by phosphorylating proteins such as FtsZ, DivIVA, MacP, and LocZ(27–30). Although the septal localization of Pkn1 resembles that of StkP, its recruitment mechanism is likely distinct because (i) Pkn1 lacks a transmembrane domain, and (ii) *C. trachomatis* lacks FtsZ, instead employing FtsK as a central organizer of its polarized cell division machinery(4).

The conserved ATP-binding residue K42 was also crucial for proper Pkn1 localization and function. Unlike Pkn1_6xH, the Pkn1(K42G)_6xH mutant rarely localized to the septum and instead exhibited diffuse or punctate localization (Fig. 1B & C). Moreover, ectopically expressed Pkn1(K42G)_6xH failed to complement the enlarged cell morphology and EB progeny production defect of *pkn1*KD (Fig. 3E-G). These findings demonstrate that K42 is essential for Pkn1 function and are consistent with a requirement for ATP binding and kinase activity in both septal localization and regulation of cell division.

Several additional findings connect Pkn1 to the regulation of septal PG assembly. Conditional knockdown of *pkn1* produced enlarged, irregular rounded organisms resembling the aberrant forms generated by β-lactam mediated inhibition of PG synthesis (Fig. 2A and S3). In addition, *pkn1*KD also reduced PG synthesis as observed by EDA-DA labeling and disrupted the septal PG ring formation, whereas ectopic expression of Pkn1_6xH during knockdown restored both PG synthesis and ring organization (Fig. 4). These data suggest that Pkn1 is critical for proper synthesis, incorporation, or spatial organization of PG at the division septum. Because metabolic labeling alone cannot distinguish among these possibilities, the precise step in PG biogenesis controlled by Pkn1 remains to be determined. To evaluate potential interactions with divisome components, we performed a BACTH analysis, which detected an interaction between Pkn1 and MreB (Fig. 5A). In support of this identified interaction, disruption of MreB polymerization with A22 caused Pkn1 to lose its septal localization and become diffusely distributed within the chlamydial cytoplasm (Fig. 5C and D). Nevertheless, because both the BACTH assay and A22 treatment have experimental limitations, additional studies will be necessary to determine whether Pkn1 binds directly to MreB in chlamydial cells, whether MreB is a substrate of Pkn1, and what domains mediate this association. Together, these observations support a model in which polymerized MreB is required to recruit or stabilize Pkn1 at the division septum, where Pkn1 phosphorylates one or more unidentified substrates that regulate septal PG assembly/organization and cell division (Fig. 6).

**Figure 6.**
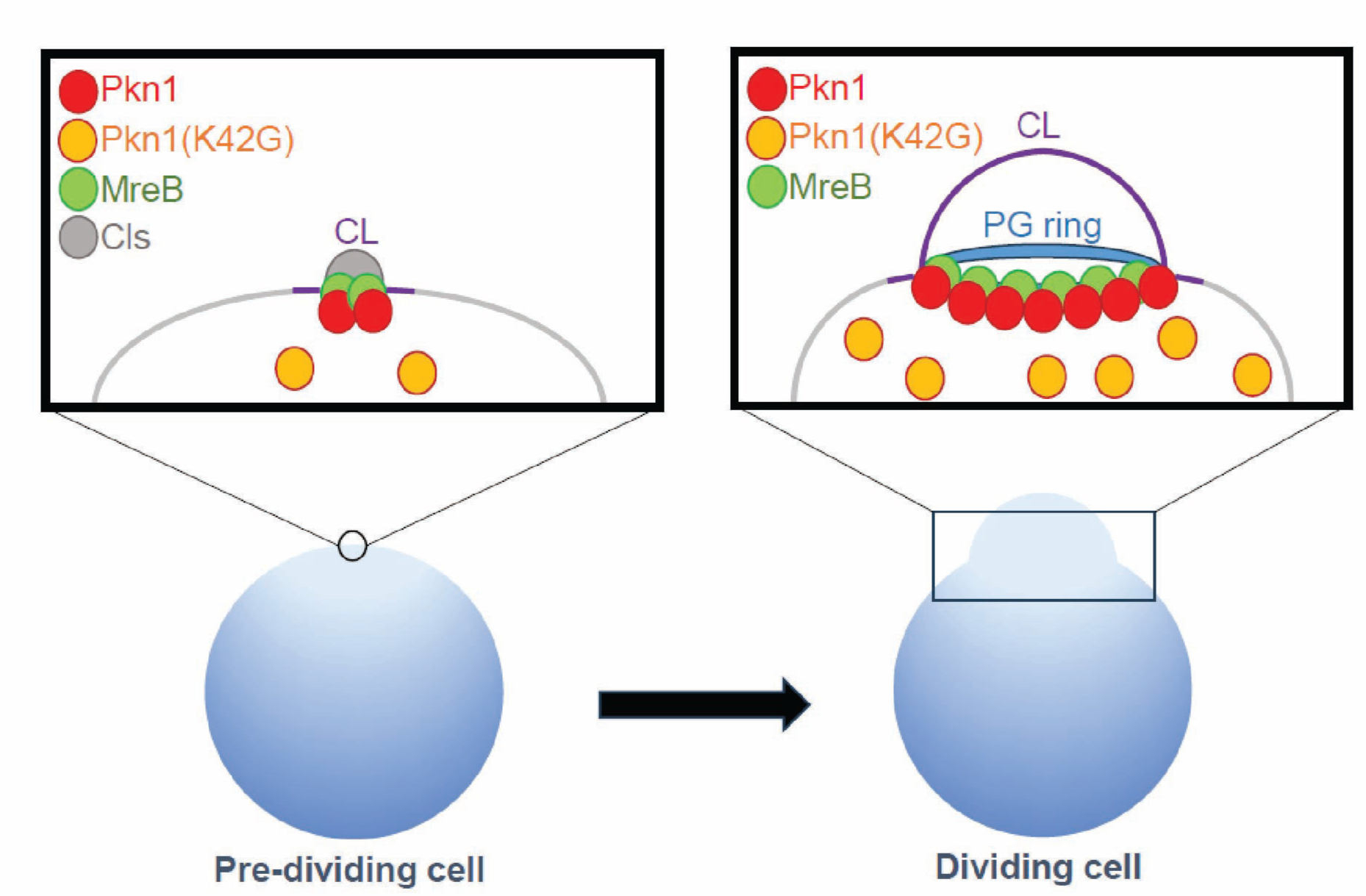
The scheme of Pkn1 function. In the pre-dividing cells, Cls is localized at the spot, which will start budding, and synthesizes cardiolipin. The cardiolipin (CL, purple) forms the positive curvature, and MreB is recruited to the site(7), and Pkn1 is also recruited to the site. In dividing cells, Pkn1 is localized at the septum forming a ring structure. MreB polymerization at the septum is crucial for this localization, and Pkn1’s kinase activity is also required for the localization.

The complementation experiments further distinguish the contributions of Pkn1 and DnlJ to the observed phenotypes. CRISPR-dCas12 targeting the *pkn1* promoter reduced the expression of both *pkn1* and the downstream gene *dnlJ* (Fig. 2D). Ectopic expression of DnlJ alone did not restore normal cell morphology or EB progeny production, whereas Pkn1 fully rescued the morphology and PG defects. These findings demonstrate that the cell division phenotype resulted primarily from depletion of Pkn1 rather than from the polar effect on *dnlJ*. In contrast, expression of Pkn1 alone partially restored EB progeny production, and complete recovery was observed only when both Pkn1 and DnlJ were expressed (Fig. S5). These results indicate that appropriate expression of both proteins is necessary for efficient EB progeny production. Because chlamydial genome levels were not substantially reduced following knockdown, the progeny defect is consistent with impaired secondary differentiation or EB infectivity rather than a general failure of genome replication. Recent studies, including work from our group, have identified several major factors that regulate chlamydial secondary differentiation(31). We demonstrated that cyclic di-AMP above a critical threshold promoted secondary differentiation, whereas reduction of cyclic di-AMP below its normal level inhibited this transition(32). In addition, the redox status of the bacteria also regulates secondary differentiation(33). Regulated protein turnover mediated by Clp proteolytic system has also been shown to contribute to control of this developmental transition(34). Although these findings have revealed several important regulatory mechanisms, we hypothesize that additional factors might be involved, and based on the results of the present study, we expect that DnlJ serves an important function during chlamydial secondary differentiation.

Many questions remain regarding the role of phosphorylation in chlamydial physiology. In particular, the direct substrates of Pkn1 and the upstream signals that control its activity have yet to be identified. Future phosphoproteomic and biochemical studies will be necessary to define the Pkn1-dependent phosphorylation network, validate direct substrates, and determine how phosphorylation alters the activities or localization of division proteins. It will also be important to establish how Pkn1 activity is coordinated temporally with MreB assembly, septal PG synthesis/organization, and secondary differentiation. Overall, our findings identify Pkn1 as an important regulator of chlamydial cell division and EB progeny production. Defining this *Chlamydia*-specific regulatory pathway may provide broader insight into the unusual physiology of this obligate intracellular pathogen and reveal potential vulnerabilities that could ultimately be exploited for the development of new therapeutic strategies against chlamydial infectious diseases.

## Materials and Methods

### Organisms and Cell Culture

McCoy (kind gift of Dr. Harlan Caldwell, NIH), mouse fibroblast-derived cell line was cultured in Dulbecco’s Modified Eagle Medium (DMEM; Invitrogen, Waltham, MA) containing 10% fetal bovine serum (FBS; Hyclone, Logan, UT) and 10 μg/mL gentamicin (Gibco, Waltham, MA) at 37°C with 5% CO_2_. *C. trachomatis* serovar L2 (434/Bu) lacking the endogenous plasmid (-pL2; kind gift of Dr. Ian Clarke, Univ. Southampton) was used for transformation. All cell cultures and chlamydial stocks were routinely tested for Mycoplasma contamination using the Mycoplasma PCR detection kit (Sigma, St. Louis, MO). For *E. coli*, NEB10β competent cells (New England Biolabs, Ipswich, MA) were used for the amplification of pBOMB-derivative vectors. *E. coli* was grown at 30°C in LB media with appropriate selection. All chemicals and antibiotics were obtained from Sigma unless otherwise noted.

### Cloning

The list of vectors and primers used in this study is detailed in Supplemental Table 1. Target genes were amplified by PCR with Phusion DNA polymerase (NEB) using 10 ng *C. trachomatis* L2 genomic DNA or appropriate vectors as a template. Some DNA segments were directly synthesized as a gBlock fragment (Integrated DNA Technologies, Coralville, IA). If plasmids were used as a template, then we treated the PCR product with DpnI enzyme to remove templates. The PCR products were purified using a PCR purification kit (Qiagen, Hilden, Germany). The HiFi Assembly reaction master mix (New England Biolabs (NEB), Ipswich, MA) was used following the manufacturer’s manual in conjunction with plasmids pBL_v2 linearized with EagI and KpnI, pBE_v2 linearized with EagI and KpnI, or pE12CRia_v2 (empty vector) linearized with BamHI. To make a complementation strain, we used the knockdown plasmids linearized with SalI. The linearized plasmids were also dephosphorylated with FastAP (ThermoFisher). The products of the HiFi reaction were transformed into NEB10β competent cells (NEB) and plated on LB agar with appropriate antibiotics. Plasmids were subsequently isolated using a mini-prep kit (Qiagen) and verified by plasmid digest and sequencing from individual colonies grown overnight in LB broth with appropriate antibiotic selection.

### Transformation of Chlamydia trachomatis

McCoy cells were plated in a six-well plate the day before beginning the transformation procedure. *C. trachomatis* serovar L2 without plasmid (-pL2) resuspended with Tris-CaCl_2_ buffer (10 mM Tris-Cl pH 7.5, 50 mM CaCl_2_) was incubated with 2 μg plasmid at room temperature for 30 min. During this step, McCoy cells were washed with 2 mL Hank’s Balanced Salt Solution (HBSS) media containing Ca^2+^ and Mg^2+^ (Gibco). After that, McCoy cells were infected with the transformants in 2 mL HBSS per well. The plate was centrifuged at 400 x g for 15 min at room temperature and incubated at 37°C for 15 min. The inoculum was aspirated, and 2 mL 1X DMEM containing 10% FBS and 10 μg/mL gentamicin was added per well. At 8 hours post infection (hpi), 1 μg/mL cycloheximide and 500 µg/ml spectinomycin were added, and the plate was incubated at 37°C until 48 hpi. At 48 hpi, the transformants were harvested and infected onto a new McCoy cell monolayer. These harvest and infection steps were repeated every 44-48 hpi until fluorescent, penicillin-resistant inclusions were observed.

### (RT-)qPCR

McCoy cells were infected with chlamydial transformants at an MOI of 0.5. At 10 hpi, 2 or 10 nM anhydrotetracycline (aTc) and 0.5 mM Theophylline were added or not to the culture medium. Total RNA and DNA were harvested at this time point from duplicate wells not treated with aTc. At 14 and 24 hpi, total RNA and DNA were collected using Trizol (Invitrogen) and DNeasy Tissue (Qiagen) kit, respectively, as described elsewhere(35). After DNase treatment of total RNA, cDNA was synthesized using Superscript III reverse transcriptase (Invitrogen). After diluting the cDNA 10-fold, 5 μl of the diluted cDNA was used as a template for qPCR. Equal masses of genomic DNA were used from each of the samples to quantify chlamydial genomes, which were used to normalize transcript data as described(36). For both cDNA and gDNA samples, qPCR reactions were prepared using 2X SYBR Green (ThermoFisher) in a total volume of 20 µl per well. Standard cycling conditions were used with a melting curve analysis to verify products. Transcripts and genome copies were assessed from at least three biological replicates.

### Indirect immunofluorescence assay (IFA)

McCoy cells were infected with chlamydial transformants as above. At 10 hpi, 2 or 10 nM aTc and 0.5 mM Theophylline were added or not, and the infected cells were fixed with fixing solution (3.2% formaldehyde and 0.022% glutaraldehyde in 1X DPBS) for 2 min and permeabilized with 90% MeOH for 1 min at 10.5 hpi or 24 hpi. The fixed cells were labeled with primary antibodies including rabbit anti-six histidine tag (Abcam, Cambridge, UK), goat anti-major outer-membrane protein (MOMP; Meridian, Memphis, TN), and mouse anti-AsCpf1 (Sigma-Aldrich, St. Louis, MO). To visualize the primary antibodies, donkey anti-goat antibody (488), donkey anti-mouse (647), or donkey anti-rabbit antibody (594) were used as secondary antibodies. The secondary antibodies were obtained from Invitrogen or Jackson Immunology (West Grove, PA). Coverslips were observed using a Zeiss AxioImager.Z2 with Apotome2 as noted in the figure legends.

### Inclusion forming unit (IFU) measurement

McCoy cells were infected with chlamydial transformants as above. At 10 hpi, 2 nM aTc and 0.5 mM Theophylline were added or not to the culture medium. At the indicated times, infected cells were harvested in 1 ml 2SP media then froze at -80°C. After thawing the lysates, the samples were serially diluted at 1:10 and used to infect McCoy cells seeded in 24-well plates. At 24 hpi, the number of GFP expressing inclusions were counted from 30 fields of view to calculate the IFUs from the original sample. Three biological replicates were performed.

### Labeling of chlamydial peptidoglycan

McCoy cells were infected with chlamydial transformants. The constructs were induced at 10 hpi, and 0.5 mM EDA-DA was added at 20 hpi. The infected monolayer cells were fixed at 24 hpi with 100% MeOH for 5 min and permeabilized with 0.5% Triton X-100 for 5 min. The samples were incubated with 3% bovine serum albumin (BSA) for 1 h. Afterward, we carried out click reaction with Molecular Probes Click iT Cell Reaction Buffer Kit (Invitrogen, Waltham, MA) as previously described(37).

### Bacterial Adenylate Cyclase Two Hybrid (BACTH) assay

The pKT25 and pUT18C vectors expressing the genes of interest or empty vectors were co-transformed into competent DHT1 cells, an adenylate cyclase-deficient strain, and spread onto M63 minimal medium plates containing 50 mg/ml ampicillin, 25 mg/ml kanamycin, 0.5 mM isopropyl-β-D-thiogalactopyranoside (IPTG), 0.2% maltose, 40 mg/ml 5-bromo-4-chloro-3-indolyl-β -D-galactopyranoside (X-Gal), and 0.04% Casamino Acids. The plates were incubated at 30°C for up to 7 days. Blue colonies indicate positive interactions, whereas no growth or small white colonies indicate no interactions.

## Statistical Analysis

To analyze the statistical significance between uninduced and induced samples of qPCR, cell diameters, and IFU data, we used two-tailed paired Student’s t-test.

## Data and materials availability

All data are available in the main text or the supplementary materials.

## Acknowledgements

This study was supported in part by an NIH/NIAID award (R21AI190577) to JL, an NIH/NIGMS award (R35GM151971) to SPO, and an NIH/NIAID award (R01AI179688) to JVC. The authors would like to thank Dr. Derek Fisher (Southern Illinois University) for the anti-Pkn1 antibody, Dr. Harlan Caldwell (NIAID/NIH) for McCoy cells, and Dr. Ian Clarke (University of Southampton) for the plasmidless *C. trachomatis* serovar L2 strain.

## Author contributions

Junghoon Lee: Conceptualization, Methodology, Investigation, Visualization, Writing – original draft

John V. Cox: Conceptualization, Writing – review and editing

Scot P. Ouellette: Conceptualization, Methodology, Visualization, Supervision, and Writing - review and editing

## Figure Legends

**Figure S1. The phenotype of Pkn1_6xH and Pkn1(K42G)_6xH at 24 hpi. A.** Pkn1 localization at the first dividing cells. Transformant encoding Pkn1_6xH was infected into McCoy cells, and the construct was induced at 4 hpi. The infected monolayer was fixed at 10.5 hpi with fixing solution containing 3.2% formaldenyde and 0.022% glutaraldehyde in 1X PBS for 2 min. And the samples were permeabilized with 90% MeOH for 1 min. MOMP and Pkn1_6xH were stained by indirect immunofluorescence assay (IFA) with Rabbit anti-six histidine and Goat anti-MOMP. **B.** IFA images of WTL2 at 24 hpi. The arrowheads represent the ring or filament structures of endogenous Pkn1. **(C-D)** McCoy cells were infected with chlamydial transformant encoding Pkn1_6xH or Pkn1(K42G)_6xH, and the constructs were induced at 10 hpi. IFA and IFU samples were prepared at 24 hpi. **C.** IFA images of Pkn1_6xH and Pkn1(K42G)_6xH at 24 hpi. The arrowhead represents the septum localized Pkn1_6xH. **D.** IFU of Pkn1_6xH. **and** Pkn1(K42G)_6xH. IFA images were acquired on a Zeiss AxioImager.Z2 equipped with an Apotome2 using a 100X lens objective. Scale bar: 1(A) and 2(B and C) µm.

**Figure S2. Quantification of the cell size of uninduced and induced samples of *pkn1*KD.** We measured the cell diameter of *pkn1*KD uninduced and induced cells through Fiji software. 129 uninduced cells and 117 induced cells were measured. UI = uninduced (i.e., -aTc); I = induced (i.e., +aTc) for all sample types. **: p<0.001 via two sample equal variance t-test.

**Figure S3. The morphology of pkn1KD is similar with the morphology treated with β-lactam antibiotics. A.** McCoy cells were infected with *C. trachomatis* transformed with an aTc-inducible plasmid encoding Pkn1_6xH. At 10 hpi, expression of the construct was induced, and the 1U of Penicillin G was treated or not. The infected monolayer was fixed with 100% MeOH at 24 hpi for 10 minutes, and the major outer membrane protein (MOMP) was stained with goat anti-MOMP antibody and the donkey anti-Goat (488). **B.** McCoy cells were infected with *C. trachomatis* carrying an aTc-inducible construct encoding CRISPRi system targeting *pkn1*. At 4 hpi, the construct was induced or not, and the infected cells were fixed at 24 hpi with 100% MeOH. The MOMP and dCas12 were stained with goat anti-MOMP, donkey anti-goat(488), mouse anti-dCas12, and donkey anti-mouse(594) antibodies. The images were acquired on a Zeiss AxioImager.Z2 equipped with an Apotome2 using a 100X lens objective. Scale bar: 2 μm.

**Figure S4. The phenotype of *pkn1*KD-DnlJ_6xH.** McCoy cells were infected with *C. trachomatis* transformed with an aTc-inducible plasmid encoding CRISPRi system targeting *pkn1* promoter and DnlJ_6xH (*pkn1*KD-DnlJ_6xH). At 10 hpi, expression of the construct was induced or not. DNA and RNA samples were collected at 10, 14, and 24 hpi. IFA and IFU samples were prepared at 24 hpi. **A.** IFA images of the *pkn1*KD-DnlJ_6xH strain at 24 hpi. Shown are individual panels for DnlJ_6xH and dCas12 labeling as well as the merged image with MOMP. **B.** Quantification of IFUs from uninduced and induced samples at 24 hpi. **C.** Quantification of genomic DNA by qPCR in uninduced and induced samples. **D.** Quantification of transcripts by RT-qPCR for *pkn1, dnlJ, euo, hctA, and omcB*. IFA images were acquired on a Zeiss AxioImager.Z2 equipped with an Apotome2 using a 100X lens objective. Scale bar: 2 µm. UI = uninduced (i.e., -aTc); I = induced (i.e., +aTc) for all sample types. *: p<0.05; **: p<0.001 via two-tailed paired Student’s t-test.

**Figure S5. The phenotype of *pkn1*KDop.** McCoy cells were infected with *C. trachomatis* transformed with an aTc-inducible plasmid encoding CRISPRi system targeting *pkn1* promoter, Pkn1 and DnlJ_6xH (*pkn1*KDop). At 10 hpi, expression of the construct was induced or not. DNA and RNA samples were collected at 10, 14, and 24 hpi. IFA and IFU samples were prepared at 24 hpi. **A.** IFA images of the *pkn1*KDop strain at 24 hpi. Shown are individual panels for DnlJ_6xH and dCas12 labeling as well as the merged image with MOMP. **B.** Quantification of IFUs from uninduced and induced samples at 24 hpi. **C.** Quantification of genomic DNA by qPCR in uninduced and induced samples. **D.** Quantification of transcripts by RT-qPCR for *pkn1, dnlJ, euo, hctA, and omcB*. IFA images were acquired on a Zeiss AxioImager.Z2 equipped with an Apotome2 using a 100X lens objective. Scale bar: 2 µm. UI = uninduced (i.e., -aTc); I = induced (i.e., +aTc) for all sample types. *:p<0.05 via two-tailed paired Student’s t-test

